# Microtubule retrograde flow retains neuronal polarization in a fluctuating state

**DOI:** 10.1101/2021.09.01.458567

**Authors:** Max Schelski, Frank Bradke

## Abstract

In developing vertebrate neurons, a neurite is formed by more than a hundred microtubules. While individual microtubules are dynamic, the microtubule array itself has been regarded as stationary. Using live cell imaging in combination with photoconversion techniques and pharmacological manipulations, we uncovered that the microtubule array flows retrogradely within neurites to the soma. This microtubule retrograde flow drives cycles of microtubule density, a hallmark of the fluctuating state before axon formation. Shortly after axon formation, microtubule retrograde flow slows down in the axon, which stabilizes microtubule density cycles and thereby functions as a molecular wedge to enable axon extension. We propose microtubule retrograde flow and its specific slowdown in the axon to be the long-sought mechanism to single one neurite out to drive neuronal polarization.

## RESULTS AND DISCUSSION

Microtubules support the complex morphology of neurons and form arrays of parallel tracks to transport cargo within vertebrate neuronal projections (neurites) (*1*). Within the microtubule array, single microtubules are dynamic, and can be transported (*2*), generated (*3, 4*) as well as polymerize and depolymerize (*5*). The microtubular array in neurites itself, however, is regarded as a stationary structure, even during neuronal development (*1, 6*). We challenge this view by showing that microtubules in developing neurites flow retrogradely from the distal tip to the cell body. This microtubule retrograde flow (MT-RF) then slows down in the axon to establish neuronal polarity.

To visualize the dynamics of the microtubule array in neurites, we expressed a microtubule subunit (tubulin) fused to the fluorophore mEos3.2 (*7*), which can be photoconverted with UV light from green to red fluorescence, in developing murine hippocampal neurons. By laser-induced local photoconversion of the fluorophore in neurites, we labelled small microtubule patches throughout the whole microtubule stack (**fig. S1, A and B**) and followed them (**Fig. 1A**) prior to axon formation. In contrast to what is expected from a stationary microtubule array, the patches quickly moved retrogradely in the neurites towards the soma, at a speed of 0.53 ± 0.03 µm/min (mean ± SEM; **Fig. 1, B** **and C; Movie S1**). Immobile patches were only observed in 4.7% of neurites and anterogradely moving patches in 1.8% of neurites. MT-RF occurred in the absence of neurite retraction (**fig. S2B**). To test whether the entire microtubule array in the neurite moved retrogradely, we followed multiple microtubule patches in the same neurite. To avoid light-induced stress-reaction in this neurite, the photoactivatable fluorophore Dronpa (*7*) was fused to tubulin, which needs less light than mEos3.2 to be photoactivated (**Fig. 1D**). The photoactivated microtubule patches moved retrogradely in a synchronous manner, keeping a constant distance to each other (**Fig. 1, E** **and F; Movie S2**). Thus, before axon formation, the microtubule array in a neurite flows retrogradely towards the cell body.

**Fig. 1.**
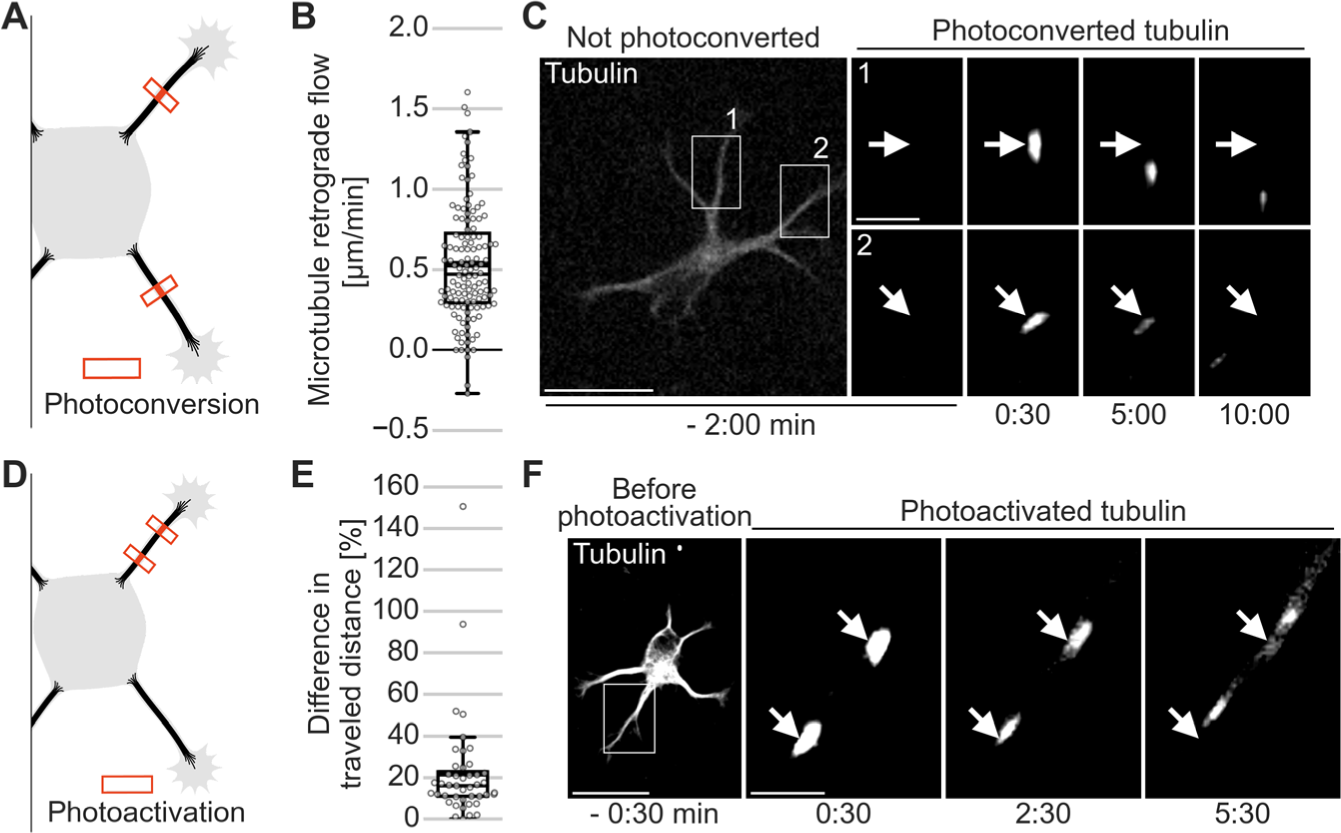
The entire microtubule array in the neurite flows retrogradely before axon formation. The tubulin subtype TUBB2a was fused either (**A - C**) to the photoconvertible fluorophore mEos3.2 or (**D - F**) to the photoactivatable fluorophore Dronpa, expressed in neurons and imaged after one day in culture. (**A**) Illustration of photoconversion experiment from panels [B and C]. (**B** and **C**) Small microtubule patches were photoconverted in two neurites of neurons without axon. (**B**) Microtubule retrograde flow in neurons without axons (n = 71 cells, N = 11 independent experiments). Each data point represents one neurite. (**C**) Representative cell for [B]. (**D**) Illustration of photoactivation experiment for [E and F]. (**E** and **F**) Two or three microtubule patches were photoactivated in a single neurite. (**E**) For each neurite, the distance difference of the patch that moved furthest and the patch that moved the least was divided by the lower distance (n = 44 cells, N = 9 independent experiments). (**F**) Representative cell for [E]. Arrows indicate location of photoconversion [C] or photoactivation [F] at 0:00 min. Thick line in boxplots shows mean. Scale bars, 20 µm overview, 5 µm for zoomed images.

Microtubules polymerize into the neurite tip to extend the neurite (*8*). As neurites remain short before neuronal polarization and after axon formation (*9, 10*) it raised the possibility that MT-RF might impede neurite growth by reducing the net extension of the microtubule array. Consequently, when the axon rapidly grows out, MT-RF may slow down in the axon to enable its accelerated growth. Upon photoconverting microtubule patches in neurites of axon-bearing neurons we found that MT-RF was similar compared to neurites of neurons without an axon. However, in the axon, MT-RF was more than two-fold slower (0.61 ± 0.08 µm/min in minor neurites compared 0.23 ± 0.05 µm/min in the axon (mean ± SEM; **Fig. 2, A** **to C; Movie S3**). We found that MT-RF was also slower in the axon when neurons were cultured in 3 dimensions (3D) (*11*) (**fig. S3, A to C; Movie S4**). Of note, at later development when neurites grow as dendrites (*12*), their MT-RF decreased but not to the level of almost stationary microtubules found in the axon (**fig. S4, A and B**).

**Fig. 2.**
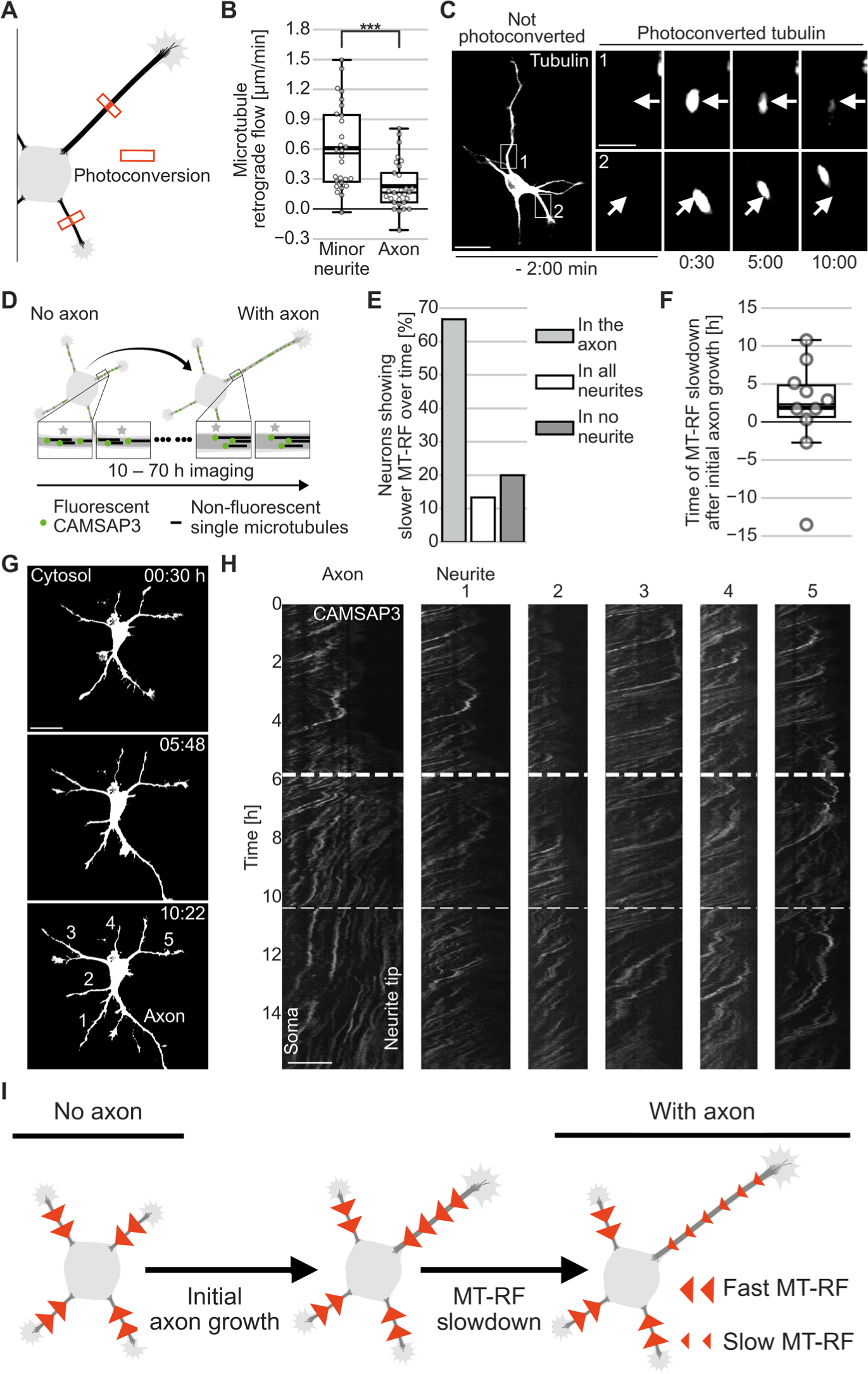
Microtubule retrograde flow slows down shortly after initial axon growth. (**A - C**) The tubulin subtype TUBB2a fused to the photoconvertible fluorophore mEos3.2 or (**D - H**) CAMSAP3 fused to the fluorophore mNeonGreen and the cytosolic fluorophore tandem-mCherry (cytosol) were expressed in neurons and imaged after one day in culture. (**A**) Illustration of photoconversion experiment from panels [B and C]. (**B** and **C**) MT-RF in minor neurites and axons (n = 28 cells, N = 8 independent experiments). Arrows indicate location of photoconversion at 0:00 min. (**D**) Illustration of long-term imaging of MT-RF by imaging the microtubule (minus) end-binding protein CAMSAP3 from panels [E – H]. (**E** and **F**) MT-RF was averaged over 200 min and was considered slowed down once it was 20% slower in the axon compared to all other neurites for at least 120 min. (n = 15 cells, from those, 10 cells for time of MT-RF slowdown, N = 6 independent experiments). (**G**) Cell morphology of neuron from [H] with the fluorophore tandem-mCherry to label the cytosol before (0:30 h) and after (5:48 h) the axon reached axon-like length and when MT-RF slowed down in the axon (10:22 h). Neurites from [H] are labelled at 10:22 h. (**H**) Neon-CAMSAP3 intensity along each neurite (x-axis) over time (y-axis; kymograph). The left site is close to the soma. Thus, a trace from top right to lower left indicates MT-RF. The thick and thin dashed line indicate the time when the future axon reached axon-like length (5:48 h) and when MT-RF slowed down in the axon compared to all other neurites (10:22 h), respectively. (**I**) Illustration of when MT-RF slows down during axon formation. Thick line in boxplots shows mean. ***P < 0.001, Dunn’s test. Scale-bar, 20 µm for overview, 5 µm for zoomed images.

We then defined when MT-RF slowed down in relation to axon formation. As repeated photoconversions are toxic for neurons, we measured MT-RF during neuronal polarization in all neurites simultaneously by expressing and imaging the minus-end marker calmodulin-regulated spectrin-associated protein 3 (CAMSAP3) (*13*) fused to a fluorophore (*14*) (**Fig. 2, D** **to H**). MT-RF slowed down in the axon compared to all other neurites in 66.7% of neurons (**Fig. 2E**). In these neurons, MT-RF decreased relative to all other neurites more than 1 h after initial axon growth (**Fig. 2, F** **to I; Movie S5**). Thus, the slowdown of MT-RF in the growing axon could act as a molecular wedge that enables the steady growth of the axon.

Before axon formation, neurons are in a fluctuating state, where each neurite transiently acquires axon-like properties through accumulating axon-specifying microtubule bound signals (*14-16*). MT-RF could be the basis of this fluctuating state by transporting these signals (*9, 16–18*) from the neurite back to the cell body. We reasoned that MT-RF could constantly deplete axon-specifying factors if it remains fast in axon-like neurites. To label axon-like neurites we used the constitutively active motor domain of kinesin 1 family proteins (caKIF5C), a key marker for axon-like properties that labels neurite tips and, after axon formation, persistently labels the tip of the axon (*14*) (**Fig. 3A**). Before axon formation, MT-RF was indistinguishable in neurites that accumulate caKIF5C compared to MT-RF in neurites that did not accumulate this marker (**Fig. 3, B** **to D; Movie S6**). By contrast, MT-RF decreased at the transition of axon formation in neurites with caKIF5C accumulation (**fig. S5, A to C; Movie S7**). Thus, persistent MT-RF slowdown is specific to the axon and does not occur in neurites with axon-like properties. This suggests that before axon formation, MT-RF constantly acts on neurites and could keep the neuron in an unpolarized state.

**Fig. 3.**
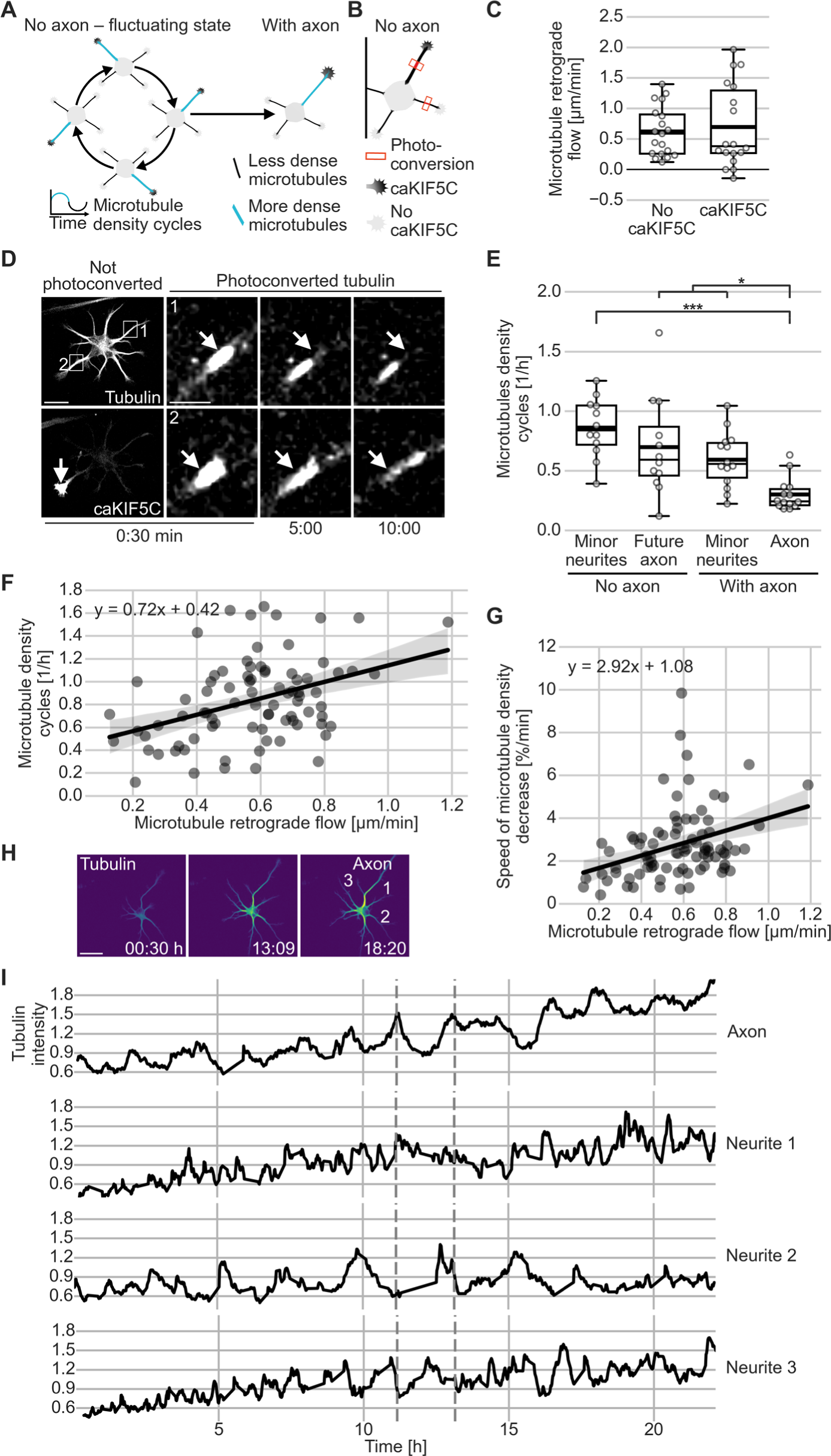
Microtubule retrograde flow keeps neurons in the fluctuating state. Neurons were cultured for one day expressing (**C** and **D**) the tubulin subtype TUBB2a fused to the photoconvertible fluorophore mEos3.2 and caKIF5C fused to the fluorophore Cerulean3 (**E** - **I**) or the fluorophore mNeonGreen fused to CAMSAP3 and TUBB2a fused to the fluorophore mScarlet. (**A**) Illustration of fluctuations of microtubule density and caKIF5C before axon formation. (**B**) Illustration of photoconversion experiments for [C and D]. (**C** and **D**) MT-RF quantified in neurites with and without caKIF5C accumulation in neurons without axons (n = 18 cells, N = 8 independent experiments). White arrows in photoconverted channel of [D] indicate areas of photoconversion and in the caKIF5C channel points to the growth cone with caKIF5C accumulation. (**E - I**) Microtubules and MT-RF were imaged simultaneously in neurons from before to after axon formation. Changes of average microtubule density in a neurite of 30% or more of the average density in the neuron was considered half a cycle. For a full cycle an increase followed by a decrease, or vice versa, was needed (n = 14 cells with axon, of those, 12 cells had data prior to axon formation, N = 3 independent experiments). (**E**) Microtubule density cycles in neurons before and after axon formation. Microtubule density cycles were averaged for all minor neurites of one neuron. (**F** and **G**) Correlation and linear regression of MT-RF (F) with microtubule density cycles (Pearson r = 0.39, p = 0.0002) and (G) with the speed of microtubule density decrease during cycles (Pearson r = 0.35, p = 0.001) of single neurites. The formula for the linear regression line is shown in upper left of panel with the light grey area indicating the 95% confidence intervals. Each data point represents one neurite (**H** and **I**) Representative cell for [E]. (**H**) at start of imaging (0:30 h), directly after (13:09 h) and some hours after (18:20 h) axon growth. Neurite numbers from [I] are annotated at the last timeframe. (**I**) Tubulin intensity was normalized for the neuron and smoothened with a rolling window of 2 in each neurite. The first dashed line indicates indicates the end of the time period a neuron was considered to be without an axon (until 120 min before axon growth), the second dashed line indicates the time of axon formation. Thick line in boxplots shows mean. ***P < 0.001, **P <0.01, *P< 0.05, Kruskal Wallis multiple comparison with Dunn’s posthoc test with Holm-Bonferroni correction. Scale bar, 20 µm for overview, 5 µm for zoomed images.

Next, we checked whether MT-RF could deplete microtubule-bound axon-specifying markers by investigating its association with microtubule density changes in the fluctuating state. During fluctuations, microtubules cycle between low and high density in different neurites (**Fig. 3A**). While higher density is caused by transient increases in polymerization (*19*), the mechanism underlying the decrease of microtubule density remains elusive. We hypothesized that MT-RF could decrease microtubule density through moving microtubules out of the neurite to fuel microtubule density cycles. Consistent with this hypothesis, axons showed more than 2-fold fewer microtubule density cycles than minor neurites (**Fig. 3E, H** **and I; Movie S7**). Additionally, we found more microtubule density cycles in neurites with faster MT-RT prior to axon formation (Pearson r=0.39, **Fig. 3F**). Importantly, neurites with faster MT-RF reduced microtubule density faster (Pearson r=0.35, **Fig. 3G**).

This raised the possibility that fast MT-RF could prevent neurites from becoming axons by constantly counteracting stable microtubule accumulation. Consistent with this view, pharmacological manipulations that transform non-growing neurites into multiple growing axons per cell (*17, 20, 21*) slowed down MT-RF in neurites. Even before growth, taxol and para-amino blebbistatin (pa-Blebb) treatments decreased MT-RF to the level of control treated axons (**Fig. 4, A** **to C; Movie S8**). We therefore tested whether in addition to slower MT-RF microtubule density cycles also decreased, as we had found for the axon. Indeed, pa-Blebb and taxol reduced the number of microtubule density cycles from 1.05 ± 0.07 cycles/h to 0.45 ± 0.05 and 0.63 ± 0.6 cycles/h, respectively (mean ± SEM; **Fig. 4, D** **to F; Movie S9**). Thus, slower MT-RF could reduce microtubule density cycles and enable neurites to become axons by lifting the depletion of microtubules from neurites (**Fig. 4G**).

**Fig. 4.**
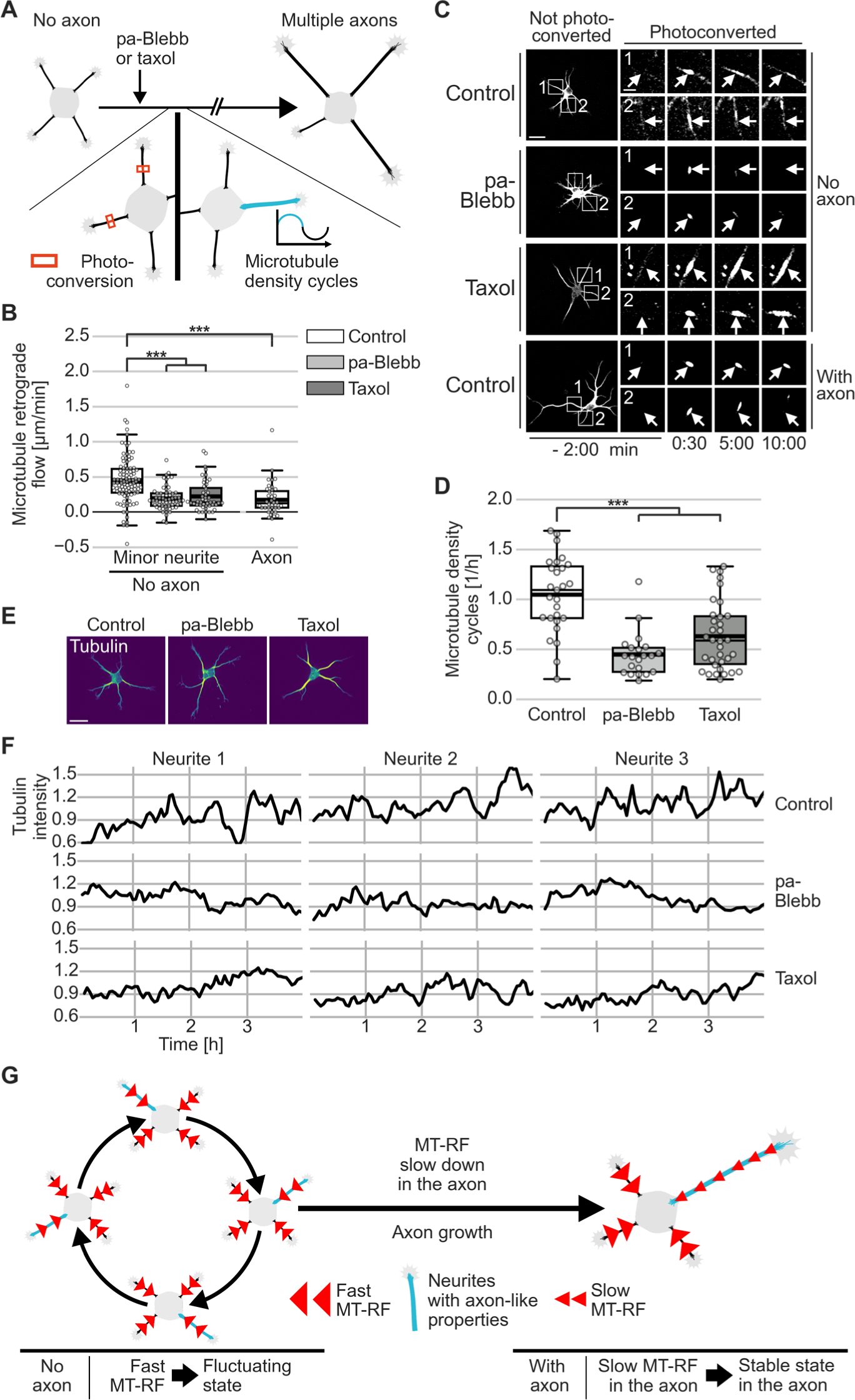
Stabilizing axon identity in multiple neurites slows down MT-RF and reduces microtubule density cycles. The tubulin subtype TUBB2a fused to either (B and C) the photoconvertible fluorophore mEos3.2 or (D–E) the fluorophore mNeonGreen was expressed in neurons, imaged after one day in culture directly after treating neurons for 20 – 240 min with para-amino blebbistatin (pa-Blebb, 40 µM), taxol (6 nM) or control (DMSO). (**A**) Illustration of photoconversion experiment after treatments of panels [B and C]. (**B** and **C**) MT-RF after pa-Blebb, taxol and control treatment. (**B**) Each data point represents one neurite. (n = 22 cells, N = 4 for taxol, 32 cells, N = 4 for pa-Blebb, 54 cells, N = 11 for control without axon and 40 cells, N = 9 for control with axon; N, number of independent experiments). Anterogradely moving patches were plotted as zero MT- RF. (**C**) Arrows indicate location of photoconversion at 0:00 min. (**D - F**) Microtubule density cycles were measured by counting the density increases followed by decreases in neurites of at least 30% of the average density in a neuron. Neurons were traced and average intensity along neurites measured using a self-made fully automated algorithm. (**D**) The number of cycles per hour were averaged for all neurites of neurons before axon formation (n = 34 cells for taxol, N = 3; 21 cells for pa-Blebb, N = 2; and 27 cells for control, N = 3; N, number of independent experiments). (**E**) Morphology and color-coded tubulin intensity (blue lower, yellow higher intensity) of neurons at start of imaging. (**F**) Tubulin intensity was normalized for each neuron and then smoothened with a rolling window of 2 frames in each neurite. (**G**) Illustration of MT-RF fueling fluctuations and then MT-RF slowing down in the axon. Thick line in boxplots shows mean. ***P < 0.001, Kruskal Wallis multiple comparison with Dunn’s posthoc test with Holm-Bonferroni correction. Scale bar, 20 µm for overview, 5 µm for zoomed images.

Here we reveal that the microtubule array in neurites is not stationary but instead constantly moves towards the soma during early development. Interestingly, a mechanically stable microtubule array might be needed for stable neurite growth since neurons do not utilize the matrix as a stable scaffold to pull on (*11*). MT-RF could mechanically destabilize the microtubule array thereby inhibiting stable growth in the fluctuating state (*12, 14*). Slowdown of MT-RF in the axon could allow the axonal microtubule array to be used as a stable scaffold from which to extent. As neuronal development relies on complex architectural changes of the cell, modulating the microtubular array through MT-RF is an attractive mechanism to drive morphological changes and offers a novel perspective on how neurons can be reconfigured during development and degenerative diseases.

## ACKNOWLEDGEMENTS

We thank B. Hilton, T.-C. Lin, E. Burnside, S. Dupraz, T. Pietralla and E. Alfadil for critically reading and discussing the manuscript, K. Fährmann for independently confirming the analysis for Fig. 1B, the Animal Research Facility, the Light Microscope Facility of the DZNE Bonn, Liane Meyn and Blanca Randel for technical assistance. This work was supported by Deutsche Forschungsgesellschaft (DFG), the International Foundation for Research in Paraplegia (IRP), and Wings for Life. M.S. was supported by the Add-On Fellowship for Interdisciplinary Sciences from the Joachim Herz Foundation and a fellowship from the International Max Planck Research School for Brain and Behavior (IMPRS-BB). F.B. is a member of the excellence cluster ImmunoSensation2, the SFBs 1089 and 1158, and is a recipient of the Roger De Spoelberch Prize.

## AUTHOR CONTRIBUTIONS

M.S. conceived the project; M.S. and F.B. designed research; M.S. performed research; M.S. analyzed the data; F.B. supervised the research; M.S. and F.B wrote the paper.

## DATA AND MATERIAL AVAILABILITY

The datasets supporting the current study have not been deposited in a public repository. The reported data are archived on file servers at the German Center for Neurodegenerative Diseases (DZNE) and are available from the corresponding author on request. All plasmids used in this study will be made available through Addgene.

## SUPPLEMENTAL INFORMATION

### MATERIALS AND METHODS

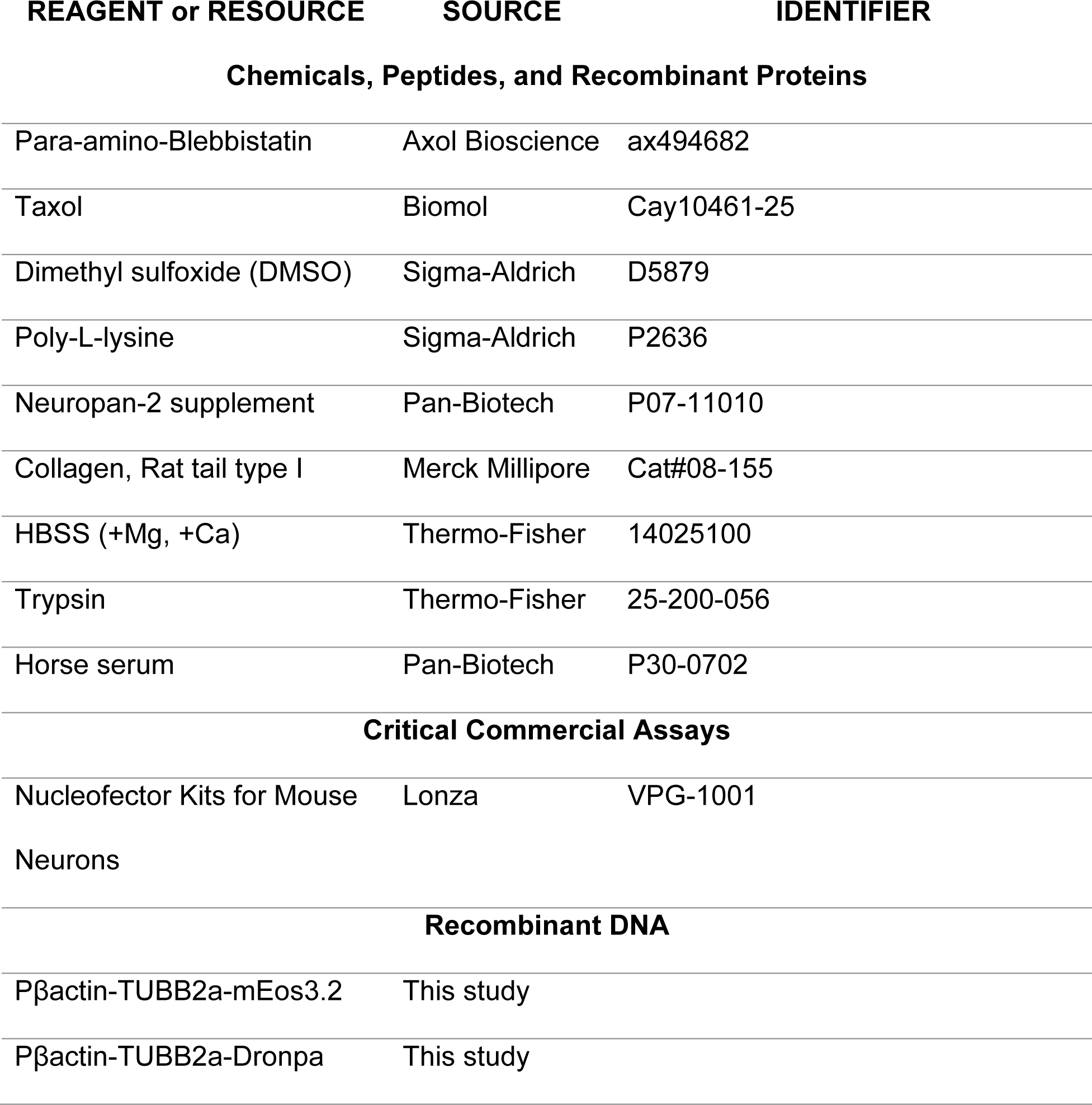
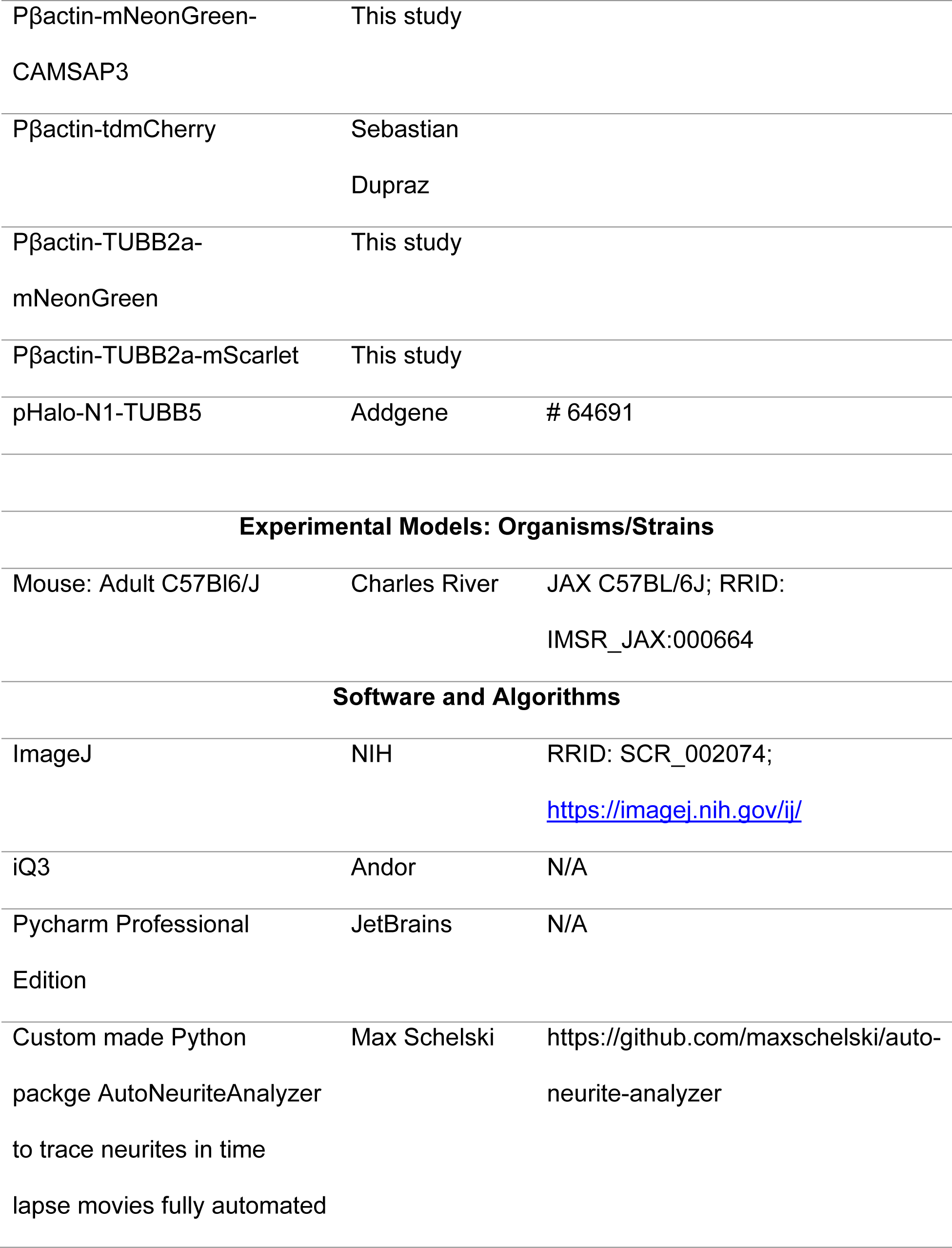
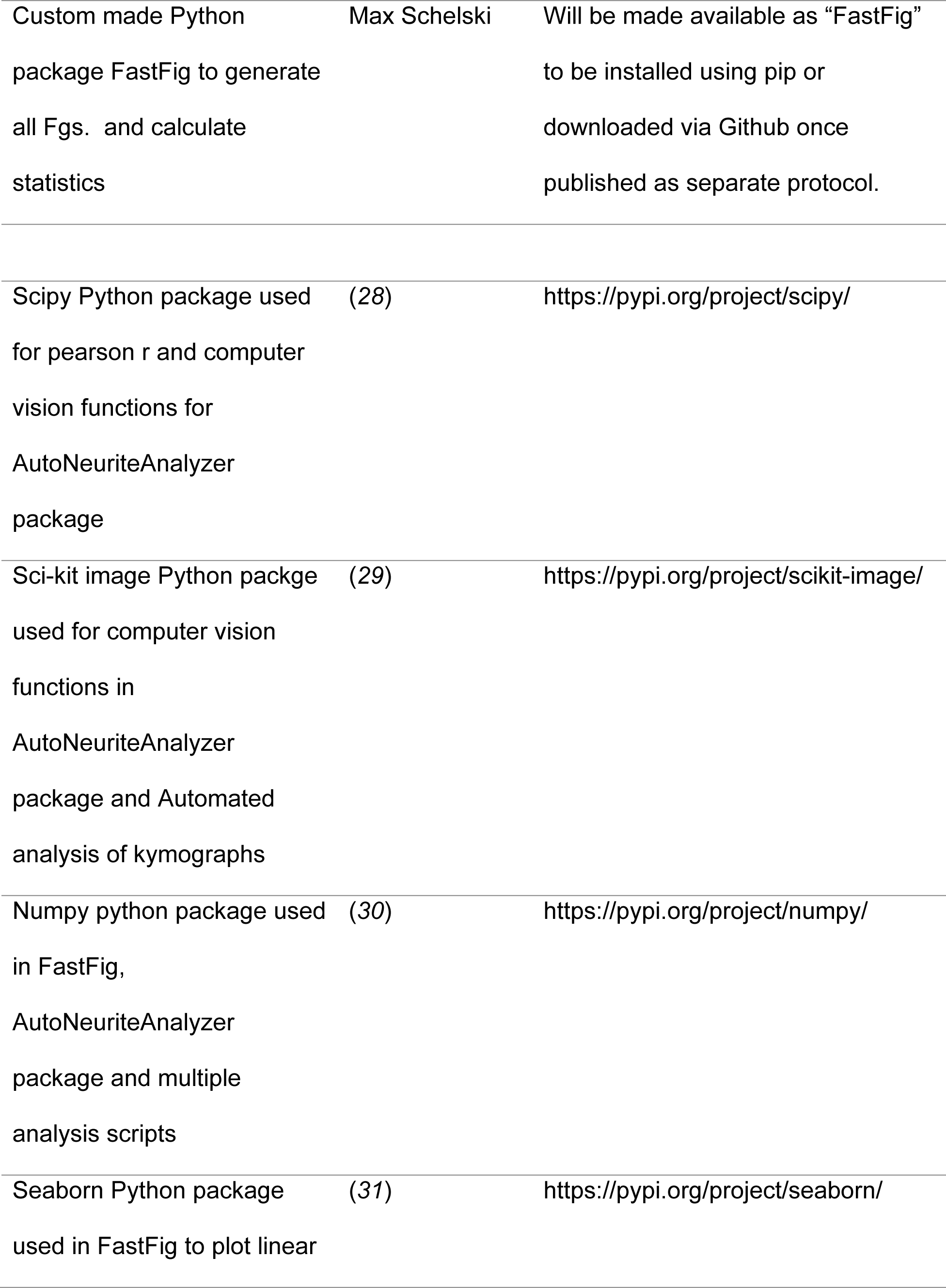
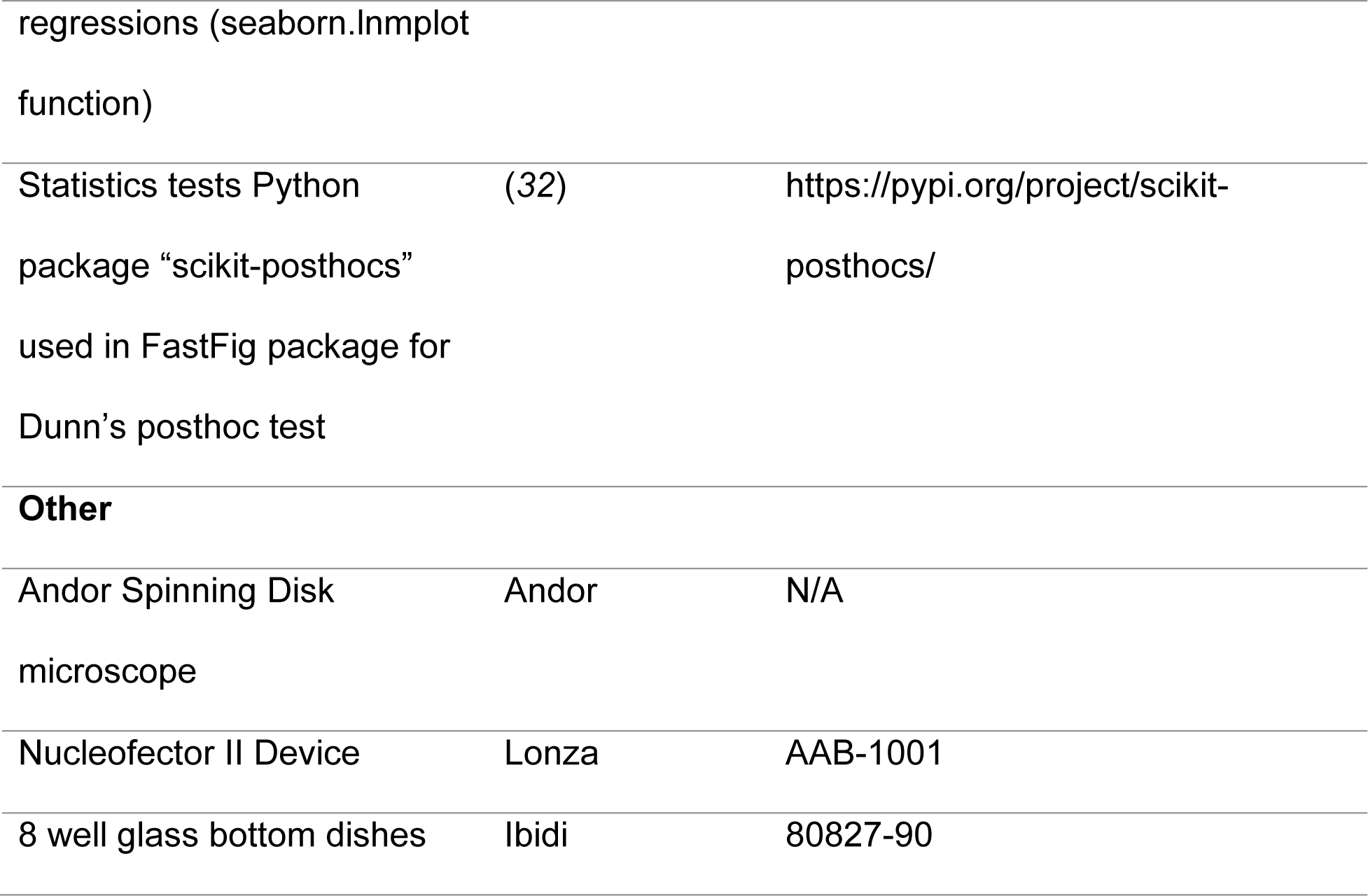
TABLE OF RESOURCES.

#### Animals

All animal experiments were performed in accordance with the Animal Welfare Act and the guidelines of the North Rhine-Westphalia State Environment Agency (Landesamt für Natur, Umwelt und Verbraucherschutz (LANUV)). Animals were group housed (up to 5 mice per cage) with room temperature controlled at 21–22°C, and an artificial 12-h light:dark cycle (lights off at 6:00 pm). Mice were given food and water *ad libitum* throughout the experiment.

#### Neuron Culture

Primary hippocampal neurons were obtained from embryonic day 16.5 (E16.5) to E17.5 mouse brains. Cells were kept at 36.5°C and 5% CO_2_. Neurons were cultured without a glia feeding layer similarly to as previously described (*23*). In brief, after dissection, hippocampi were collected in HBSS with magnesium and calcium (Thermo Fisher). Next, hippocampi were digested in 0.25% trypsin (Thermo Fisher) at 37 °C for 15 min. Hippocampi were then washed three times with MEM-1% horse serum (MEM-1% HS) media [1X MEM, 1X essential and non-essential amino acids, 2mM L-glutamine (all from Thermo Fisher), 0.22% NaHCO_3_, 0.6% glucose and 10% horse serum (Pan-Biotech)]. Lastly, hippocampi were mechanically dissociated with fire-polished glass-Pasteur pipettes. After transfection, 10 thousand or 15 thousand neurons were then plated in 8 well glass bottom dishes (Ibidi). Dishes were coated with 1mg/ml Poly-L-lysine at room temperature overnight, washed with double distilled water three to four times and then placed in the incubator at 36.5 °C and 5% CO_2_ with MEM-10% horse serum (MEM-10% HS) to equilibrate for at least 2 hours before plating neurons. Two to four hours after plating neurons, MEM-10% HS medium was replaced by neuronal media (N2) (1X MEM, sodium pyruvate 1 mM, Neuropan 2 supplement 1% (Pan-Biotech), NaHCO_3_, 0.6% glucose, 2 mM L-glutamine) which was pre-conditioned through 2-3 days incubation with a glia feeder layer.

#### Neuron transfections

Cell transfections were performed using the Nucleofector II (Lonza) with program 0-005 and the mouse neuron Nucleofector® Kit (Lonza) according to the manufacturer’s specifications. For each transfection, 5 × 10^5^ neurons were used with varying amounts for different plasmids (15 μg for Pβactin-TUBB2a-mEos3.2, 15 µg for Pβactin-TUBB2a- Dronpa, 6µg (Fig. 2) or 3 µg (Fig. 3) for Pβactin-mNeonGreen-CAMSAP3, 10 µg for Pβactin-td-mCherry, 10 µg for Pβactin-TUBB2a-mNeonGreen, 10 µg for Pβactin-TUBB2a-mScarlet and 10µ g for pHalo-N1-TUBB5). All plasmids were prepared with the EndoFree Maxiprep kit (QIAGEN).

#### Collagen 3D matrices

Neurons were grown in 3D collagen matrices as described previously (*12*). Briefly, neurons were grown in collagen matrices at a density of 0.75 to 1.5 x10^6^ cells/mL. For the collagen matrix, 225 µL collagen with a concentration of 4.13 mg/mL was combined with 30 µL 10x MEM and 15 µL 5.5% sodium bicarbonate and mixed. Transfected cells in medium were added 1:3 and mixed carefully. From the hydrogel-cell mix single 40 µL drops were added to the bottom of 8 well glass bottom dishes (Ibidi). Gels were then kept in the cell culture incubator at 37 °C for 20 min. Then around 400 µL conditioned N2 medium was added to fully cover the gels.

#### DNA constructs

All DNA plasmids made in this study were constructed in the Pβactin plasmid backbone (kind gift from C. C. Hoogenraad (University Utrecht); (*24)*). Plasmids were constructed by using homology-based cloning with the NEBuilder® HiFi DNA Assembly Master Mix (New England Biolabs, # E2621L) according to the manufacturer’s specifications. Protein coding sequences were assembled by amplifying them by PCR using Q5 polymerase (New England Biolabs, #M0491S) using primers with homology sequences in overhanging ends and subsequent clean up from an agarose gel. The plasmid backbone for these assemblies was constructed by digestion of Pβactin plasmid with NheI and HindIII and subsequent clean up from an agarose gel. One sequence was designed as homology site for the upstream sequence in the backbone (CTCTGTTCCTCCGCAGCCCCCAAGCTAGC) one for the downstream sequence (AAGCTTCGAATTCTGCAGTCGACGGTACC) and one to combine two protein coding sequences together (Linker; GGAAGTTCAGGAGGTTCTAGT, will be translated into the neutral linker sequence GSSGGSS). The same homology sequences were used for all assemblies. The protein coding sequence of TUBB2a was amplified from mouse cDNA with the upstream backbone sequence in the forward primer and the Linker sequence in the reverse primer (forward primer: ATGCGCGAGATCGTGCAC, reverse primer: TCAAGCCTCATCTTCACCCTCC). mEos3.2 was amplified from the plasmid mEos3.2-N1 (gift from Michael Davidson, Addgene #54525; http://n2t.net/addgene:64691; RRID: Addgene_64691) (*7*) using forward primer ATGAGTGCGATTAAGCCAGAC and reverse primer TCTAGATCCGGTGGATCC. Dronpa was amplified from the plasmid Dronpa-Tubulin-6 (gift from Michael Davidson, Addgene #57301, http://n2t.net/addgene:57301; RRID: Addgene_57301) using forward primer ATGGTGAGTGTGATTAAACCAGACATG and reverse primer TTACTTGGCCTGCCTCGG. CAMSAP3 was amplified from the plasmid pFasBac+GFP-CAMSAP3 (gift from Ron Vale (Addgene plasmid # 59038; http://n2t.net/addgene:59038; RRID: Addgene_59038) (*25*). MNeonGreen was amplified from the pNCS-mNeonGreen plasmid (Allele Biotechnology) using forward primer ATGGTGAGCAAGGGCGAG and reverse primer GGCCATGGTTCCGGAC. MScarlet was amplified from the plasmid pLifeAct_mScarlet_N1 (gift from Dorus Gadella; Addgene plasmid # 85054; http://n2t.net/addgene:85054; RRID: Addgene_85054) (26) using forward primer TGGTGAGCAAGGGCGAG and reverse primer CTTGTACAGCTCGTCCATGC. pHalo-N1-TUBB5-Halo was a kind gift from Yasushi Okada (Addgene plasmid # 64691; http://n2t.net/addgene:64691; RRID: Addgene_64691) (27). For fusing mEos3.2, Dronpa, mNeonGreen or mScarlet C-terminally to TUBB2a, the upstream backbone sequence was added to the forward TUBB2a primer, the reverse complemented linker sequence was added to the reverse TUBB2a primer, while for mEos3.2, Dronpa, mNeonGreen and mScarlet the linker sequence was added to forward primers and the reverse complement of the downstream backbone sequence was added to reverse primers. To fuse mNeonGreen N-terminally to CAMSAP3, for mNeonGreen, the upstream backbone sequence was added to the forward and the reverse complement of the linker sequence to the reverse primer while for CAMSAP3 the linker sequence was added to the forward and the reverse complement of the downstream backbone sequence to the reverse primer.

#### Live cell imaging

All imaging experiments were performed on an Andor spinning disk Nikon Eclipse Ti microscope with Perfect Focus system (Nikon) and a Yokogawa CSU-X1 Spinning Disk Unit (CSUX1-A1N-E/FB2 5000rpm Control, FW, DMB 95L100016) using a Plan Apo VC 60x Water objective N.A. 1.2 (**fig. S3**, Nikon) or Plan Apo VC 60x Oil objective N.A. 1.4 with an additional 1.5 magnifier (**fig. S1**) or no additional magnifier (all other Fgs., Nikon) and iQ3 software (Andor). A dichroic Quad filter (for wavelengths: 395-410; 482-492; 561-568; 628-650) and a single bandpass emission filter for the green channel (525/50; Semrock Brightline), the red channel (617/73; Semrock Brightline) and the far-red channel (697/75; Semrock Brightline) were used. For most photoconversion experiments (**Figs. 1, B and C; 2, B and C; 3, C and D; 4, B and C, and figs. S2 to S5**) a quad band emission filter was used (440/40, 521/21, 607/34, 700/45; Semrock Quad Band Filter FF01-440/521/607/700) to allow faster imaging without changing filters. For the green, red and far-red channel a 488 nm laser (100 mW), a 561 nm laser (100 mW) and a 640 nm laser (100 mW) were used, respectively (REVOLUTION 500 series AOTF Laser modulator and combiner unit, solid state laser modules). Experiments were conducted inside a large incubator chamber around the microscope, kept at 37 °C. A small chamber for 8-well ibidi glass bottom dishes (Pecon, #000470) was supplied with humidified air mixed with CO_2_ to a concentration of 6-7% CO_2_. Images were taken with an Andor iXON camera (DU897) in CCD mode at 1 MHz horizontal readout rate and with a pixel size of 0.44 µm (2 x 2 binning; **Figs. 1, B and C; 2, B and C; 3, C and D; 4, B and C, and figs. S2 to S5**), 0.22 µm (no binning; **Figs. 1, E and F; 2, E-H; 3, E-I; 4, D-F**) or 0.15 µm (no binning, 1.5x magnifier; **fig. S1**).

Images were taken every 60 seconds (**Figs. 2** **E to H, and 3 E to I**), every 3 min (**Fig. 4 D** **to F**) or as specific in the section “Photoconversion and photoactivation”.

For all experiments neurons were selected based on a representative morphology of more than 2 (usually 4 or more) neurites. Imaging was done under lowest possible light exposure. Usually long exposure times (800 - 1000 ms) and lower laser power were preferred over higher laser power and short exposure times. For example, for long term imaging experiments of up to 70 h (**Fig. 2, E** **to H**) neurons were imaged with 488 nm at 4% and 800 ms exposure time and with 561 nm at 5% and 200 ms exposure time.

For drug treatments (**Fig. 4**) 40 µM pa-Blebb, 6 nM taxol or DMSO as control (all to a final DMSO concentration of 0.2%) were added to cells after one day in culture for 20 to 240 min directly prior to the experiment.

#### Photoconversion and photoactivation

For photoconversion and photoactivation experiments the Fluorescence Recovery After Photobleaching and Photo Activation (FRAPPA) illumination system was used (Andor, Revolution). The FRAPPA module focuses the laser on single pixels for a specified time (dwell time) and moves through all pixels for a specified number of repetitions. The area for photoconversion was chosen to be small (specific sizes for experiments described below) to keep phototoxicity minimal.

In experiments with photoconversion of microtubule patches (**Figs. 1, B and C; 2, B and C; 3, C and D; 4, B and C, and figs. S2 to S5**), small microtubule patches were photoconverted in two neurites. Photoconversion was performed with the 405 nm laser (0.37 µW power, 400 repetitions, 20 µs dwell time, 2 pixel x 1 pixel or 1 pixel x 1 pixel area, 0.44 µm pixel size). After photoconversion, images were acquired in the photoconverted (red) and not photoconverted (green) channel every 30 s for 20 min (**fig. S4**) or 10 minutes (**Figs. 1, B and C; 2, B and C; 3, C and D; 4, B and C, and figs. S2 to S4**). For most experiments in **Figs. 1, B and C; 2, B and C**, images of all cells were taken again 1-3 h after photoconversion to check whether cells survived the experiment. Cells which died were excluded from analysis.

In experiments with photoconversion of microtubule patches in neurons grown in 3D collagen matrices (**fig. S3**) photoconversion was done with the 405 nm laser (0.5 µW power, 400 repetitions, 20 µs dwell time, 2 pixel x 1 pixel area, 0.44 µm pixel size). After photoconversion, images were taken every 2 min for 10 min. Z-stacks were acquired at a z-step size from 1.7 µm to 1.85 µm. Compared to experiments in dishes on PLL (**Figs. 1, B and C; 2, B and C**), we observed retraction of neurites more often during the experiment. To exclude an effect of retraction on measured MT-RF, we excluded neurons that showed noticeable neurite retraction. To check whether neurites retracted, the entire neuron was imaged with a larger z-stack following the experiment.

To test whether photoconversion reaches the entire microtubule stack (**fig. S1**), all neurites of a neuron were photoconverted with the 405 nm laser (0.37 µW power, 400 repetitions, 20 µs dwell time, 2 pixel x 1 pixel area, 0.44 µm pixel size). Z-stacks were acquired with a z step-size of 0.25 to 0.26 µm.

In experiments where microtubule patches were photoactivated (**Fig. 1 E** **and F**) two or three microtubule patches were photoactivated in a single neurite of a neuron. Before photoactivation, the entire cell was bleached by taking a brief 200 ms image with high power (50%) of the 488 nm laser. Photoactivation was done with the 405 nm laser with less than 1% of total light intensity required for photoconversion of mEos3.2 (0.04 µW power, 40 repetitions, 20 µs dwell time, 4 pixel x 1 pixel area, 0.22 µm pixel size). After photoactivation, one image was taken every 1 min for 6 min. During imaging, Dronpa gradually recovered fluorescence. To prevent background intensity from going too high, before every acquired frame, a part of the soma was bleached with the 488 nm laser (3.51 µW power, 10 repetitions, 20 µs dwell time, 10 pixel x 12 pixel area, 0.22 µm pixel size).

#### Analysis of photoconversion in whole microtubule stack

For **fig. S1** chromatic shift between the red (TUBB2a-mEos3.2, photoconverted tubulin) and the far-red channel (TUBB5-Halo, whole tubulin) needed to be corrected for. Intensity in the red channel was higher than in the far-red channel further away from the cover glass. Chromatic shift was determined by measuring which z-slice number had the highest intensity in each channel and then averaging the difference between these z-slice numbers across all neurites (around 1.8 z-slices, which was rounded to 2 z-slices). Average intensity for all neurites in both channels was measured in ImageJ after manually thresholding both channels similarly, to only exclude neurite background. Intensity of the highest z-slice in the red channel was then divided by the intensity in the far-red channel 2 z-slices below that (to account for chromatic shift).

#### Registering low-intensity time-lapse movies

Time-lapse movies, which were used to generate kymographs (**Fgs. 2, E to H; 3, D to I; 4, D to F**), were registered through correcting for translation shifts using a self-made Python script. The script was developed since standard ImageJ registering plugins (e.g., StackReg) did not yield satisfying results, likely due to low intensity of registered channels. To register a time-lapse movie, correlation of a reference image with the frame to be registered was calculated at up to one pixel shift for each dimension for 8 shifts in total. If correlation improved in one direction, the grid was moved by one pixel in that direction and correlations were calculated again, until correlation did not improve in any direction. Best shift values in x and y were used as starting shift values for the next frame to account for gradual shifts in time-lapse movies. Images were shifted by rolling them in shift direction (numpy.roll). For reference images, it was important to roll and not to move images by adding zero values, since zero values lead to problems when registering low intensity images. The reference image was changed every 70 frames to the newest shifted images to account for changes in morphology over time. Registered images were saved with zero values instead of rolled area in shifted areas to make the shift clearly visible.

#### Kymograph generation

Time-lapse movies from which kymographs were generated were registered with a self-made Python script (section “Registering low-intensity time-lapse movies”). Kymographs were generated manually in ImageJ using the KymoResliceWide plugin (https://imagej.net/plugins/kymoreslicewide) and multi-line tool with a line width of 11 pixel and maximum intensity extracted across that width. Lines were always oriented so that the left part of kymographs corresponded to the direction of the soma and the right part to the direction of the neurite tip.

#### Automated analysis of neurite length from kymographs

For [**Fig. 3, E** to **G**, **and fig. S1**] length of neurites was automatically analyzed from kymographs using a self-made Python script. In these kymographs rows from top to bottom corresponded to increasing timepoints in the time-lapse movie while columns from left to right corresponded to points along the neurite from soma to neurite tip. Lines for kymographs were always at least 5 pixels longer than the maximum neurite length, to cover the background. First, kymographs were smoothened (scipy.ndimage.median_filter, size=2), then thresholded by using the third highest value for the three pixels furthest away from the soma in all timepoints as threshold. In the thresholded kymograph, the first and last timepoint were always added to the thresholded area to allow subsequent filling of binary holes (scikit-image.morphology.binary_fill_holes). The resulting thresholded area only included the neurite. Neurite length was determined for each timepoint as distance from the soma to the pixel furthest from the soma that still belongs to the neurite (one pixel before the first not thresholded pixel closest to the soma).

#### Classification of axon formation

For all experiments, neurons were classified as not having an axon when the longest neurite was shorter than 30 µm or less than 5 µm longer than the second longest neurites and shorter than 50 µm. Neurons were classified as at the axon transition (**Fig. S5**) if the longest neurite was at least 30 µm long and at least 5 µm longer than the second longest neurite. Neurons were classified as having an axon if the longest neurite was at least 50 µm long and 10 µm longer than the second longest neurite.

#### Analysis of microtubule retrograde flow

To measure MT-RF in 2D after tubulin photoconversion (**Fgs. 1, B and C; 2, B and C; 3, B and C, and figs. S2, S4, S5**), movement of microtubule patches was traced. The mid-point of microtubule patches was estimated by eye and movement along the neurite over time was traced in ImageJ using the multi-line tool. Only microtubule patches which could be traced for more than 1 minute were analyzed.

To analyze MT-RF in neurons without axons together with neurite growth (**fig. S2**) the maximum intensity along neurites (x-axis) was plotted against time (y-axis; kymograph). Image data from **Fig. 1, B** **to C** was used. Using the ImageJ line tool, the slope of the moving photoconverted microtubule patch in the kymograph was measured by dividing the width (distance) through the height (time). Length was analyzed automatically from kymographs generated from the not-photoconverted channel as described in the section above (“Automated analysis of neurite growth from kymographs”). For **Fig. 1, B** **to C** one neurite was excluded from analysis (speed of 3.7 µm/min) since its speed was 70% higher than the second highest value and more than 8 standard deviations away from the mean.

To analyze movement of multiple microtubule patches in the same neurite (**Fig. 1, D** **to F**), Movement of two to three microtubule patches in the same neurite were traced, using the same procedure as for photoconverted microtubule patches. For all combinations of up to three patches, the difference in traveled distance was calculated. The highest difference was divided by the lower of the two traveled distances to obtain the relative difference in traveled distance.

To analyze MT-RF in neurons cultured in 3D collagen matrices (**fig. S3**), movement of microtubule patches across x, y and z was traced. For that, the mid-point of patches was located by eye and marked using the ImageJ point-tool for different timepoints in the z-slice with the highest intensity of the microtubule patch. These points were used to calculate the distance the microtubule patch moved.

For measuring MT-RF with fluorescently labeled CAMSAP3 (**Figs. 2, D to H, and 3, E to I**) kymographs were generated (see section “Kymograph generation”). In kymographs, CAMSAP3 was traced by using the ImageJ line tool. Speed was calculated as the width (distance) divided by the height (time) of the line, whereas the angle was used to calculate direction of CAMSAP3 signals. For all analysis only retrogradely moving CAMSAP3 signals were considered. For each timepoint, MT-RF speed was calculated as median speed of all traces. To analyze slowdown of MT-RF in the axon (**Fig. 2, D** **to H**), MT-RF was smoothened with a window size of 200 min. This allowed us to exclude brief moments when MT-RF in another neurite slowed down to the level of the axon. MT-RF was determined to have slowed down once smoothened MT-RF was at least 20% slower in the axon compared to all other neurites for at least 180 min.

#### Fully automated script for neurite analysis (AutoNeuriteAnalyzer package)

We developed a Python package (AutoNeuriteAnalyzer) that traced and analyzed neurites in time-lapse movies. The package worked in multiple stages for each time frame: first it found the soma of the neuron, then it found the ideal threshold for the neuron, after that, neurites were skeletonized and traced. The last step was the analysis of intensities within neurites. Briefly, the soma was extracted by using a grey opening algorithm (scipy.ndimage.grey_opening), edge extraction (scikit-image.filters.scharr), automated thresholding (scikit-image.filters.threshold_otsu) and then filling the soma from these edges (scipy.ndimage.binary_fill_holes). To obtain the threshold for the neuron, first the lowest threshold value was found that extracts the soma with the same size as the edge analysis. Then the threshold was incrementally increased, the thresholded image was skeletonized (scikit-image.ndimage.skeletonize) to get a rough neurite skeleton. The final threshold was found at the step before 5 of more pixels of the neurite skeleton were lost. Then gaps in neurites in the thresholded image, which might have been introduced by thresholding, were closed by connecting segments detached from the soma again with the soma. This was done by finding the point closest to the soma for each detached segment and connecting it to the closest point connected to the soma. Only connections were allowed which went in the approximately same direction as detached segments, which prevented these segments from being connected to a wrong nearby neurite, which could be closer. The dilated thresholded soma (scikit-image.morphology.binary_dilation) was removed from the resulting image, followed by skeletonization to obtain neurite skeletons. For each neurite, branch points were identified, which separated different sub-branches of a neurite, and then all branches of each neurite were constructed, with each branch starting from the soma. Terminal sub-branches shorter than 1 µm and neurites with less than 40 points (9 – 15 µm) were excluded. Neurites with different starting points but overlapping in different timeframes were excluded to prevent wrongly traced neurites from being analyzed. In addition, different branches that overlapped 70% or more in different timeframes were excluded. This led to some neurites not being traced or some neurites not traced for the entire movie. Tracing of neurites was most problematic when a neurite was crossed by another neurite since crossing points sometimes were interpreted as branching points. By only using branch points when intensities of emerging branches were within a similar range, the mix-up was reduced. The script only worked well with markers that label the cytosol or neurites and not when mostly the neurite tip was labeled (e.g., for Actin or F-Actin). The package is available on github (https://github.com/maxschelski/auto-neurite-analyzer).

#### Analyzing microtubule density cycles

For **Fig. 3, E** **to I**, tubulin intensity was obtained from kymographs (see section “Kymograph generation”) of the tubulin channel. To obtain average intensities of tubulin in neurites, kymographs were thresholded using the maximum background value manually determined in that channel. For each timepoint average intensity in the thresholded area was calculated. Instead of using kymographs, for **Fig. 4, D** **to F**, intensities were measured by a self-made fully automated Python script (see section “Fully automated script for neurite tracing” above). For each neurite only the branch present in most timeframes was analyzed. Tubulin intensities were then used (**Fgs. 3, E to I, and 4, D to F**) to analyze microtubule density cycles using a self-made Python script. To analyze the frequency of microtubule density cycles (**Fgs. 3E, and 4, D to F**), intensity values were normalized to the average intensity in each neuron across all timepoints. Then intensity traces were smoothened using an averaging rolling window with a window size of 2. Next, cycles were determined as changes of at least 30% (0.3) of the average intensity in the neuron across all timepoints. One cycle consisted of an increase of 0.3 followed by a decrease of 0.3 or vice versa. The number of these cycles was divided by the total time to obtain the frequency of cycles. For **Fig. 3E** the frequency was averaged for all minor neurites of the neuron. For **Fig. 4D** neurons with an axon were excluded and the frequency of microtubule density cycles was averaged for all neurites of each neuron. For **Fig. 3G** the speed of the decrease in microtubule density was calculated as the speed of tubulin intensity decrease from the highest intensity within one cycle to the following lowest intensity. All decrease speeds for one neurite were then averaged. For **Fig. 3, F** **and G** neurons were only considered before a neurite reached axon-like length. To correlate MT-RF with microtubule density cycles (**Fig. 3, F** **and G**) Pearson r was calculated using the python package SciPy (scipy.stats.pearsonr). Linear regression lines were plotted using the function lmplot from the Python package seaborn (seaborn https://pypi.org/project/seaborn/).

For **Fig. 3, E** **to I** the timepoint of axon formation was determined by using neurite length automatically measured from kymographs (section “Automated analysis of neurite length from kymographs”). As before, an axon was defined as being at least 50 µm long and at least 10 µm longer than the second longest neurite. Neurons were determined as not having an axon until 120 min before axon formation.

#### Statistics

For all data where more than two groups were compared, a Kruskal Wallis test (scipy.stats kruskal_wallis) was done first. If the resulting p-value was below 0.05, then all combinations of groups were compared. All comparisons were done (also when comparing two groups) using Dunn’s test (scikit-posthocs.posthoc_dunn), with Holm correction (step-down method using Bonferroni adjustments).

#### Python scripts

All scripts were written using Python 3.7 or 3.8 with environments build with conda (Anaconda Inc.; 2020. Available from: https://docs.anaconda.com/) and using the following packages: numpy 1.20.3 (https://pypi.org/project/numpy/), matplotlib 3.34 or 3.4.2 (https://pypi.org/project/matplotlib/), pandas 1.3.0 (for FastFig) or 1.1.3 (https://pandas.pydata.org/), SciPy 1.6.2 (for FastFig) or 1.5.2 (https://pypi.org/project/scipy/), seaborn 0.11.1 (for FastFig) or 0.11.0 (https://pypi.org/project/seaborn/), scikit-posthocs 0.6.7 (https://pypi.org/project/scikit-posthocs/) and python-pptx 0.6.19 (https://pypi.org/project/python-pptx/).

#### Figure and movie preparation (FastFig)

We developed Python package (FastFig) to prepare all Fgs. and movies. The Python package allowed for a highly standardized generation of publication-grade Fgs. and movies using simple Python scripts. The package will be published as a separate protocol and will be made available via github and installation via pip. Among the features are simple data plotting (data is automatically plotted, statistics are calculated and then annotated in the plot, and more) simple display of representative cells (automated calculation of which cells are closest to the mean, images extraction from ImageJ Hyperstacsk, defining zoom regions, annotate channels or timepoints, and more), automated generation of movies with the same scripts and flexibility as generating Fgs. (additionally, title pages, showing specific frames longer, repeating the movie, and more is possible) and all panels are aligned accurately according to a user-defined grid. Illustrations were made in Adobe Illustrator and Microsoft Powerpoint, while the illustration of the 3D matrix in Fig. S3 (without neurons) is from Biorender (<u>BioRender.com</u>).

## SUPPLEMENTARY FIGURES

**Fig. S1.**
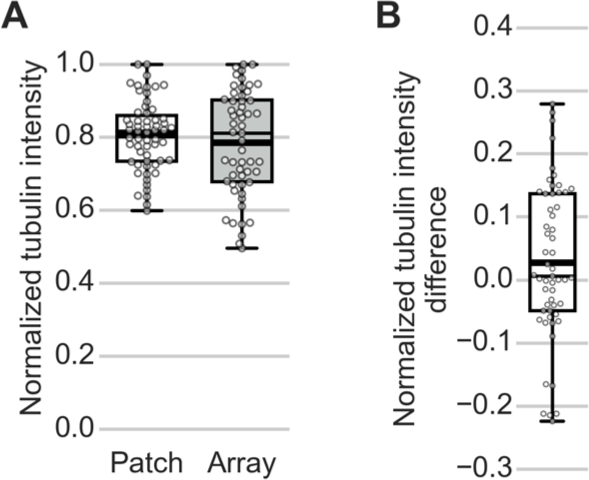
Microtubules throughout the array are photoconverted. Neurons expressing the tubulin subtype TUBB2a fused to the photoconvertible fluorophore mEos3.2 and the tubulin subtype TUBB5 fused to the halo-tag stained with a far-red chemical fluorophore (JF646, Promega) were cultured for one day and then imaged. (A and B) After photoconversion of TUBB2a-mEos, z-stacks of the entire height of all neurites were acquired. Intensities were normalized to the slice with the highest intensity in that channel. The normalized intensity at the slice furthest away from the glass was compared for the entire microtubule array and the photoconverted microtubule patch. (**A**) Comparison of the normalized intensity for the microtubule patch and the microtubule array at the topmost imaged slice of each neurite. (**B**) Difference in the normalized intensity for each neurite (55 neurites, 14 cells, N = 2 independent experiments). Thick line in boxplots shows mean.

**Fig. S2.**
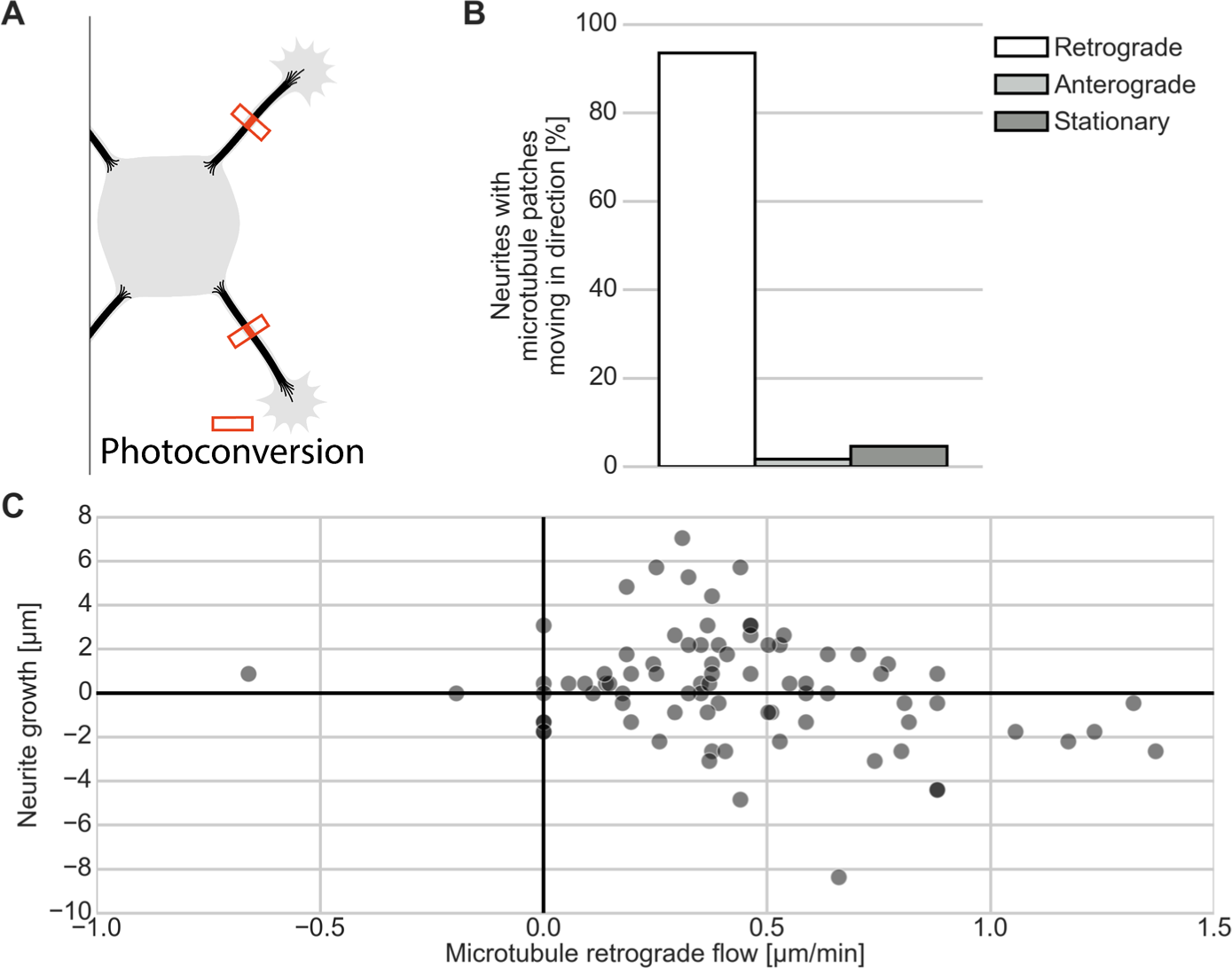
Microtubules flow retrogradely in neurons in absence of neurite retraction. Neurons expressing the tubulin subtype TUBB2a fused to the photoconvertible fluorophore mEos3.2 were cultured for one day and then imaged. (**A**) Illustration for photoconversion experiment in [B and C]. (**B**) Percentage of neurites out of all neurites imaged with microtubules moving retrogradely (towards the soma), anterogradely (towards the neurite tip) and not moving. Data from Fig. 1B (n = 71 cells, N = 11 independent experiments). (**C**) Neurite growth during 10 min imaging interval (x-axis) plotted against microtubule retrograde flow (y-axis). Part of image data from Fig. 1B reanalyzed to measure neurite growth and microtubule retrograde flow together. (n = 42 cells, N = 10 independent experiments).

**Fig. S3.**
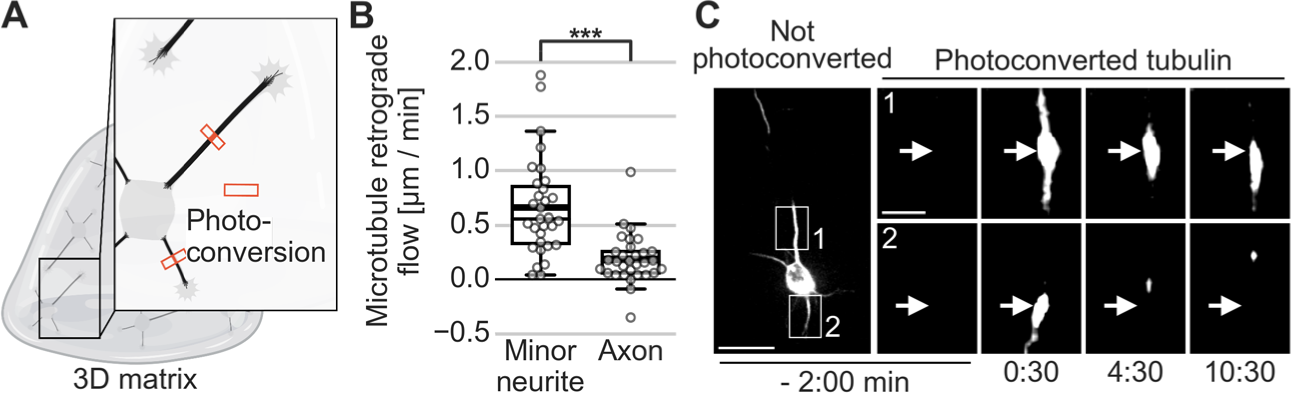
In neurons grown in 3-dimensional matrices, MT-RF slows down in the axon. The tubulin subtype TUBB2a was fused to the photoconvertible fluorophore mEos3.2, expressed in neurons and then imaged in neurons after one day in culture. (**A**) Illustration of photoconversion experiment for [B and C]. (**B**) MT-RF of neurons with axon grown in 3D. Patches that moved anterogradely were plotted with zero MT-RF. Thick line in boxplots shows mean. (**C**) Maximum intensity projection of representative cell from [B] with photoconversion done at 0:00 min and white arrows pointing to the site of photoconversion (n = 31 cells, N = 4 independent experiments). ***P < 0.001, Dunn’s test. Scale bar, 20 µm for overview images, 5 µm for zoomed images.

**Fig. S4.**
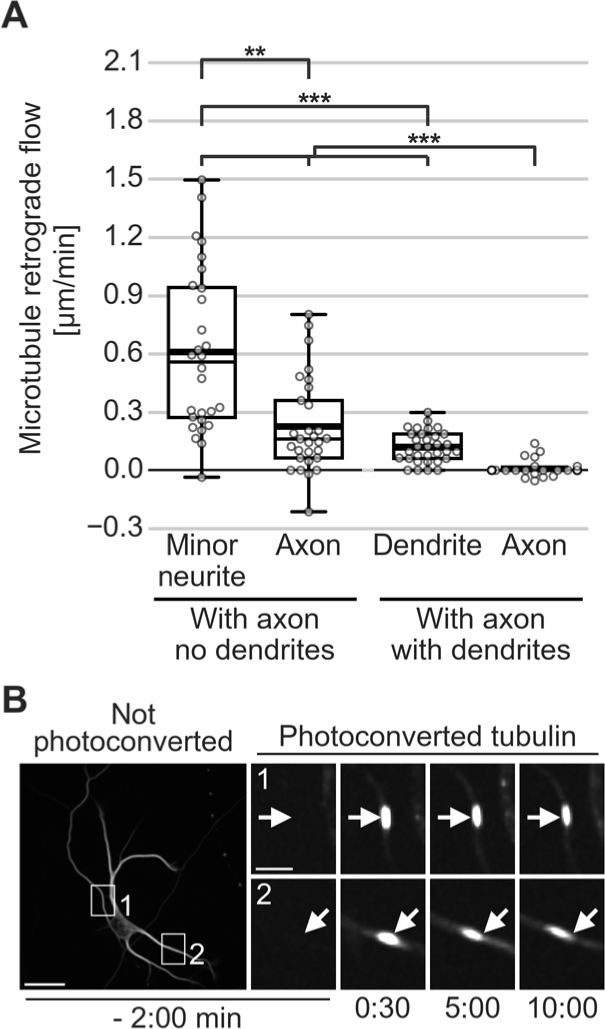
Later in development MT-RF slows down in dendrites and in the axon. The tubulin subtype TUBB2a was fused to the photoconvertible fluorophore mEos3.2, expressed in neurons and then imaged in neurons after 1 day (“With axon, no dendrites”) or after 6 to 7 days (“With axon, with dendrites”). (**A**) Quantification of MT-RF. Microtubule patches that moved anterogradely were plotted as zero MT-RF. Data from condition “With axon, no dendrites” the same as in [Fig. 2B]. (n = 26 neurons, N = 8 independent experiments for neurons without dendrites from Fig. 2B; and n = 32 neurons, N = 2 independent experiments for neurons with dendrites). Thick line in boxplots shows mean. (**B**) Representative neuron with dendrites for [A] with photoconversion done at 0:00 min and white arrows pointing to the site of photoconversion. ***P < 0.001, **P <0.01, *P< 0.05, Kruskal Wallis multiple comparison with Dunn’s posthoc test with Holm-Bonferroni correction. Scale bar, 20 µm for overview images, 5 µm for zoomed images.

**Fig. S5.**
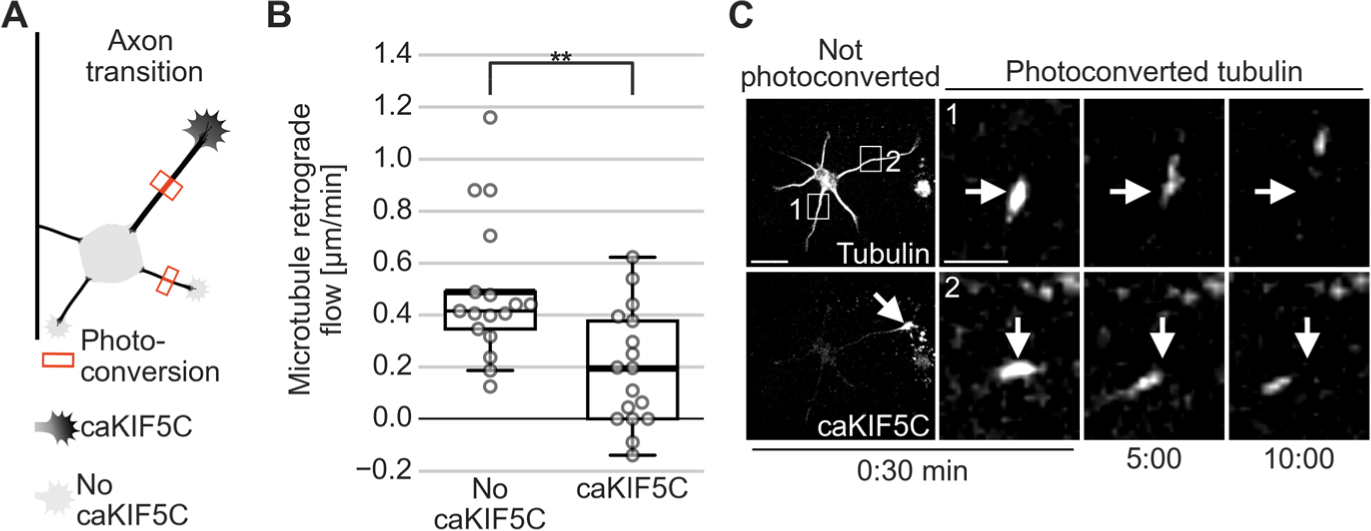
MT-RF slows down in neurites with caKIF5C at the axon transition. The tubulin subtype TUBB2a was fused to the photoconvertible fluorophore mEos3.2 and caKIF5C fused to the fluorophore Cerulean3, expressed in neurons and then imaged in neurons after 1 day. (**A**) Illustration of photoconversion experiments for [B and C]. (**B and C**) MT-RF in neurites with and without caKIF5C accumulation in neurons at the transition of growing the axon. Neurons where the longest neurite was at least 30 µm long and at least 5 µm longer than the second longest neurite was considered to be at the transition of growing the axon. Microtubule patches that moved anterogradely were plotted as zero MT-RF (n = 17 cells, N = 7 independent experiments). White arrows in photoconverted channel of [C] indicate areas of photoconversion and in the caKIF5C channel points to the growth cone with caKIF5C accumulation.

## SUPPLEMENTARY MOVIES

**Movie S1.**

**Microtubules move retrogradely towards the soma.** Data from Fig. 1C. Neurons expressing the tubulin subtype TUBB2a fused to the photoconvertible fluorophore mEos3.2 were cultured for one day and then imaged in neurons without axon. The white circle indicates the time of photoconversion at 0:00 min with white arrows indicating the site of photoconversion.

**Movie S2.**

**Multiple microtubule patches move retrogradely towards the soma in a synchronous manner.** Data from Fig. 1F. The tubulin subtype TUBB2a was fused to the photoactivatable fluorophore Dronpa, expressed in neurons, and imaged after one day in culture in neurons without axon. White arrows indicate the site of photoactivation at time 0:00.

**Movie S3.**

**MT-RF slows down in the axon.** Data from Fig. 2C. The tubulin subtype TUBB2a was fused to the photoconvertible fluorophore mEos3.2, expressed in neurons which were imaged after one day in culture. The white circle indicates the time of photoconversion at 0:00 min with white arrows indicating the site of photoconversion

**Movie S4.**

**MT-RF slows down in axons of neurons cultured in 3D.** Data from Fig. S3C. The tubulin subtype TUBB2a was fused to the photoconvertible fluorophore mEos3.2 and expressed in neurons. Neurons were cultured in 3D collagen matrices and imaged after one day in culture. Maximum intensity projection is shown. The white circle indicates the time of photoconversion at 0:00 min with white arrows indicating the site of photoconversion

**Movie S5.**

**MT-RF slows down shortly after axon outgrowth.** Data from Fig. 2, G and H. The microtubule minus-end binding protein CAMSAP3 was fused to the fluorophore mNeonGreen and expressed together with the cytosolic marker td-mCherry (cytosol). The fusion proteins were expressed in neurons and imaged after one day in culture.

**Movie S6.**

**MT-RF does not continuously slow-down before axon formation in neurites with axon-like properties.** Data from Fig. 3D. The marker for axon-like properties caKIF5C was fused to the fluorophore Cerulean3 together with the tubulin subtype TUBB2a fused to the photoconvertible fluorophore mEos3.2. Neurons without axon were imaged after one day in culture. White arrows in the photoconverted and not photoconverted channel indicate the site that was photoconverted at time 0:00. The white arrow in the caKIF5C channel points to the neurite with caKIF5C accumulation.

**Movie S7.**

**MT-RF slows-down in neurons at the transition of growing their axon in neurites with axon-like properties.** Data from Fig. S5. The marker for axon-like properties caKIF5C was fused to the fluorophore Cerulean3 together with the tubulin subtype TUBB2a fused to the photoconvertible fluorophore mEos3.2. Neurons at the beginning of growing an axon (axon transition) were imaged after one day in culture. White arrows in the photoconverted and not photoconverted channel point to the site that was photoconverted at 0:00 min. The white arrow in the caKIF5C indicates the neurite with caKIF5C accumulation.

**Movie S8.**

**Microtubule density cycles are reduced in the axon.** Data from Fig. 3, H and I. The tubulin subtype TUBB2a was fused to the fluorophore mScarlet and expressed in neurons. Neurons were imaged after one day in culture. The white arrow appears at the time when the axon grows out and points to the neurite that becomes the axon.

**Movie S9.**

**Treatment with compounds that stabilize multiple axon-identities slow down MT-RF.** Data from Fig. 4C. The tubulin subtype TUBB2a was fused to the photoconvertible fluorophore mEos3.2. and expressed in neurons. Neurons were cultured for one day and then treated with control (DMSO), 40 µM pa-Blebb or 6 nM taxol for 20 – 240 min and then directly imaged. The white circle indicates the time of photoconversion at 0:00 min with white arrows indicating the site of photoconversion

**Movie S10.**

**Treatment with pa-Blebb and taxol reduces microtubule density cycles.** Data from Fig. 4, E and F. The tubulin subtype TUBB2a was fused to the fluorophore mNeonGreen and expressed in neurons. After one day in culture, neurons were treated with control (DMSO), 40 µM pa-Blebb or 6 nM taxol for 120 min and then imaged.

